# A unifying model of species abundance distribution

**DOI:** 10.1101/2024.06.14.599104

**Authors:** Yingnan Gao, Ahmed Abdullah, Martin Wu

## Abstract

The species abundance distribution (SAD) is one of the most fundamental and best-studied macroecological patterns at the core of any biodiversity theory. Remarkably, almost every community investigated to date shows a hollow curve, indicative of the presence of many rare species and a few abundant species. While the precise nature of SAD is believed to reflect fundamental ecological processes underlying community assembly, ecologists have yet to identify a single model that comprehensively explains all SADs. Recent studies using large datasets suggested that logseries best describes animal and plant communities^1,2^ while lognormal is the best model for microbes^3^, thereby challenging the notion of a unifying SAD model across the tree of life. Using a large dataset of ∼30,000 globally distributed communities spanning animals, plants and microbes from diverse environments, here we show that powerbend distribution, predicted by a maximum information entropy-based theory of ecology, emerges as a unifying model that accurately captures SADs of all life forms, habitats and abundance scales, supporting the existence of universal ecological principles. Our findings reject the notion of pure neutrality and support the idea that community assembly is driven by both random fluctuations and deterministic mechanisms, such as interspecific trait variation and resource competition. We also show that the previously estimated one trillion microbial species existing on Earth might be orders of magnitude off.

## Main

The species abundance distribution follows one of ecology’s oldest and most universal laws. It describes the commonness and rarity in ecosystems, namely, the abundance (number of individuals) of each species in a community. Remarkably, almost every animal or plant community investigated to date is dominated by a few species, and most species in the communities are rare^4^. This universal hollow curve of SAD holds true in communities of different spatial scales, habitat types, and taxonomic groups. In recent years, such a hollow-curve pattern, known as ‘rare biosphere’ to microbiologists^5^, is also found to be universal in microbial communities.

The universality of the hollow curve SADs is both surprising and informative. It suggests that there might be universal principles operating across habitats, taxonomic groups, and spatial scales. In particular, the shape of SAD is thought to reflect key ecological processes involved in community assembly. If we can explain this high degree of unevenness, then we can gain insight into the mechanisms that structure communities, whether they involve stochastic processes, deterministic processes (e.g., species traits and niche partitioning), or a combination of both. As a result, SAD has been extensively studied in animals and plants, and dozens of models have been proposed to explain the shape of the SAD (see^4^ for a review). Among them, the best-known models are logseries^6^, lognormal^7^, broken-stick^8^, geometric series^9^ and Zipf power law^10^.

These models range from purely statistical ones selected for optimal data fitting^6^ to those explicitly incorporating ecological processes^11–13^. For example, while logseries was initially developed by Fisher to fit empirical data^6^, it has subsequently been predicted by Hubbell’s neutral theory^14^ and Harte’s maximum entropy theory of ecology (METE)^15^. METE, considered one of the most successful unifying theories of biodiversity, is based on the maximum information entropy principle (MaxEnt). It posits that the most likely form of an ecological pattern is the one that represents the most unbiased or least-informative distribution given a set of ecological constraints (e.g., the average species abundance). These constraints represent the deterministic factors that are believed to be imposed on an ecosystem. This MaxEnt framework therefore offers a platform for investigating and assessing factors that underlie the universal hollow-curve nature of SADs. For example, using MaxEnt and modeling intrinsic species trait differences, the powerbend SAD model has been proposed^16,17^. Powerbend is a modified version of the power law, distinguished by its establishment of an upper limit on the abundances of the most dominant species within a community^18^. This model, exceptionally versatile in nature, encompasses all aforementioned SAD models, with the exception of the lognormal model (Supplementary Information). Nonetheless, powerbend remains relatively obscure compared to other models and has not been widely tested.

Due to the fundamental importance of SAD in biodiversity studies, there has been a long-standing quest for a unifying SAD model for all life forms^4^. The lognormal and logseries distributions are the two most successful SAD models, and they are often used as benchmarks to test other models. For example, White et al. and Baldridge et al. tested several commonly used SAD models in about 16,000 animal and plant communities from terrestrial, aquatic, and marine environments. They found that logseries is the overall best SAD model based on the Akaike Information Criterion (AIC)^1,2^. However, in another large-scale study of over 20,000 bacterial and archaeal communities, Shoemaker et al. found that lognormal, not logseries, best describes microbial SADs^3^. The finding that microorganisms and macroorganisms may have distinct SADs raises a key question: are there unifying macroecological rules and ecological theories that can explain SADs across the tree of life?

To address this question, we test whether there is a universal SAD model that unites all types of organisms, large and small. Using a large dataset encompassing animals, plants, and microbes, we establish the emergence of powerbend as a unifying SAD model across communities of broad scales, habitats and taxonomic groups. Our findings support the idea that there are universal ecological principles governing the assembly of animal, plant and microbial communities and both deterministic and neutral processes are key drivers of community assembly.

## Results

### Powerbend is the best SAD model for animal and plant communities

In this study, we tested four SAD models — lognormal, logseries, power law, and powerbend (Extended Data Table 1). The selection of the lognormal, logseries, and power law models was based on their widespread use and extensive testing in large-scale studies of animal, plant and microbial communities ^1–3^. In contrast, the more flexible powerbend model has received little attention previously. To enable direct comparisons with previous findings, we utilized the datasets compiled by Baldridge et al. and Shoemaker et al. in our study. In terms of the goodness of fit, as measured by the modified coefficient of determination 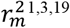, powerbend explains an average of 93.2% of the variation (weighted by the size of datasets representing different taxonomic groups, Fig. 1a) in 13,819 animal and plant SADs (Extended Data Table 2). In comparison, Poisson lognormal explains 94.7% of the variation, while logseries explains 73.2% (Fig. 1a). Using Monte-Carlo simulations, we found that powerbend, Poisson lognormal and logseries have 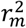 values not significantly different from 1.0 (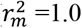 indicates a perfect fit) in 99.5%, 100% and 88.7% of SADs, respectively. Furthermore, compared to the other models, powerbend produces unbiased predictions regardless of the scale of species abundance (Fig. 1a). Poisson lognormal, while equal to powerbend in terms of the overall predictive power, tends to overestimate the abundance of the most abundant taxa (Figs. 1a and b), and performs poorly in predicting the evenness and rareness of the SAD (see below). In contrast to the other models, power law fits the data poorly 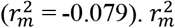 calculated using unweighted samples showed similar results (Extended Data Table 3).

**Fig 1.**
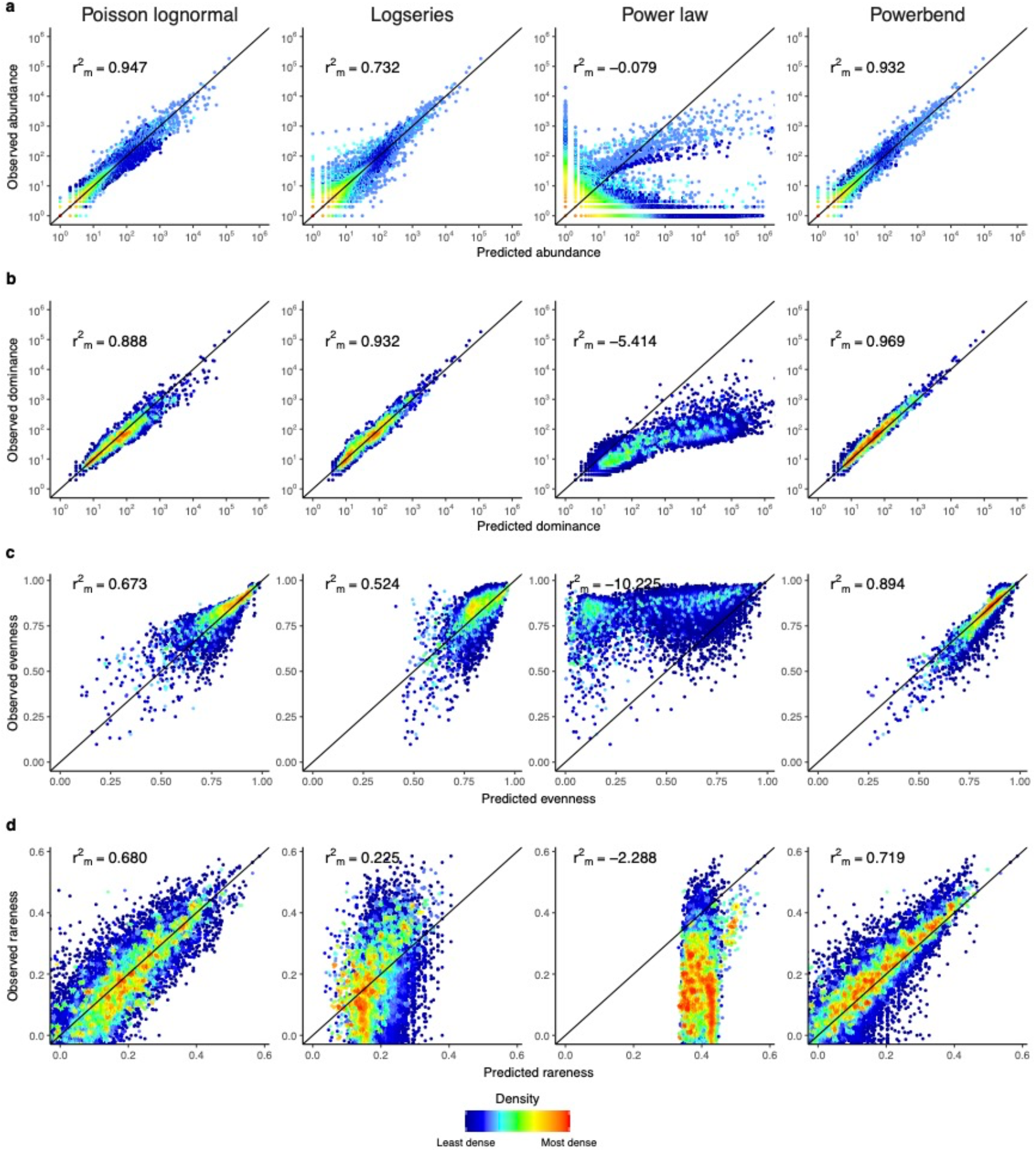
Goodness of fit of SAD models in 13,819 animal and plant communities. The predicted values are plotted against observed values for **a**, species abundance. **b**, SAD dominance. **c**, SAD evenness, and **d**, SAD rareness. Each dot in **a** represents a species and each dot in **b**-**d** represents a community. The color represents the density of the dots: red represents the densest dots and dark blue represents the least dense ones. The black diagonal line is the 1:1 line that represents a perfect fit. Goodness of fit for each SAD model is determined by the modified coefficient of determination against the 1:1 line 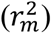 and its mean value weighted by the sample sizes of the datasets is shown. Because 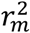 is calculated against the 1:1 line with a fixed intercept of 0, it is possible that its value drops below zero, which indicates a poor fit.

Our simulations show that when the number of observed species in a SAD is less than 40, AIC-based model selection does not have enough power to distinguish SAD models, often favoring the simpler models even when the simpler model is incorrect (Extended Data Fig.1). The small number of species in the Baldridge et al. dataset (weighted mean: 36.8 species per SAD) limited the power of AIC-based model selection in animal and plant communities. According to AIC, powerbend is significantly better than logseries in 20.88% of animal and plant SADs (*Δ*AIC≥2), while logseries is significantly better only in 0.04%. Similarly, powerbend is significantly better than Poisson lognormal in 16.44% of SADs, while Poisson lognormal is significantly better in 11.17%.

### Powerbend provides the best fit to microbial communities

Although 16S rRNA sequencing is widely employed for surveying microbial communities, there are certain challenges when using 16S rRNA sequence data to test SAD models. One key challenge arises from the fact that we only count the number of reads from a species (commonly defined as an operational taxonomic unit, or OTU, at 97% sequence identity threshold), rather than the actual number of individual cells present. To establish a connection between 16S rRNA read numbers and absolute species abundances, it is essential to account for the sampling effort within the 16S rRNA sequencing pipeline by incorporating a sampling error such as the Poisson distribution.

Shoemaker et al. have shown that the lognormal model appears to be the best SAD model for microbial communities^3^. However, it is important to note that the lognormal model was the only model in that study to incorporate a Poisson sampling error. This could confer an inherent advantage to the lognormal model over the other SAD models because 16S rRNA sequencing inherently involves multiple sampling processes. To select the best SAD model in microbial communities, we fitted Poisson lognormal and three models (logseries, power law and powerbend) with and without a Poisson sampling error to 15,329 microbial SADs (Extended Data Table 2). Incorporating a Poisson sampling error substantially improves the fit of power law and powerbend (Table 1). The powerbend model outperforms all other models in 67.0% of communities tested, while the next best model, lognormal, is superior in 18.3% of the communities (Table 1). In stark contrast, logseries, though a decent model for animal and plant SADs, is the best model in only 0.2% of microbial communities. In a direct comparison of the two best models, the Poisson powerbend model outperforms the Poisson lognormal model in 76.1% of communities, 87.0% of which are statistically significant (*Δ*AIC≥2). This finding was consistent regardless of the sequence similarity thresholds used to define microbial species (Extended Data Table 4) or whether 16S rRNA copy number variation was taken into account in determining the species abundance (Extended Data Table 5).

**Table 1.** Frequencies of each SAD model and model family being selected as the best model by AIC in 15,329 microbial communities.

| **SAD model** | **Sampling error structure** | **Best model (statistically significant)** | **Best model family (statistically significant)** |
| --- | --- | --- | --- |
| Lognormal | Poisson | 18.3% (12.7%) | 18.3% (12.7%) |
| Logseries | None | 0.2% (0.0%) | 0.2% (0.0%) |
|  | Poisson | 0.0% (0.0%) |  |
| Power Law | None | 1.7% (0.0%) | 14.5% (0.5%) |
|  | Poisson | 12.8% (0.5%) |  |
| Powerbend | None | 6.7% (2.0%) | 67.0% (56.1%) |
|  | Poisson | 60.3% (49.4%) |  |

A best model is considered statistically significant when its AIC difference to the second-best model is greater than 2.

Poisson powerbend exhibits an excellent overall fit to the observed SAD data, on average explaining 99.3% of the variation (Fig. 2a). The excellent fit is consistent throughout the entire range of species abundance spanning 7 orders of magnitude. In contrast, although Poisson lognormal and Poisson power law also provide great overall fits, they substantially overestimate the abundance of the abundant species in the communities (Fig. 2a). Using Monte-Carlo simulations, we found that Poisson powerbend exhibits 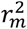 values not significantly different from 1.0 in 90.1% of SADs. This suggests that powerbend accurately describes the vast majority of microbial SADs. In contrast, Poisson lognormal and Poisson power law models have 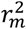 values not significantly different from 1.0 only in 67.3% and 43.0% of SADs, respectively. Poisson logseries fits the data poorly, only explaining 12.9% of variation (Fig. 2a).

**Fig 2.**
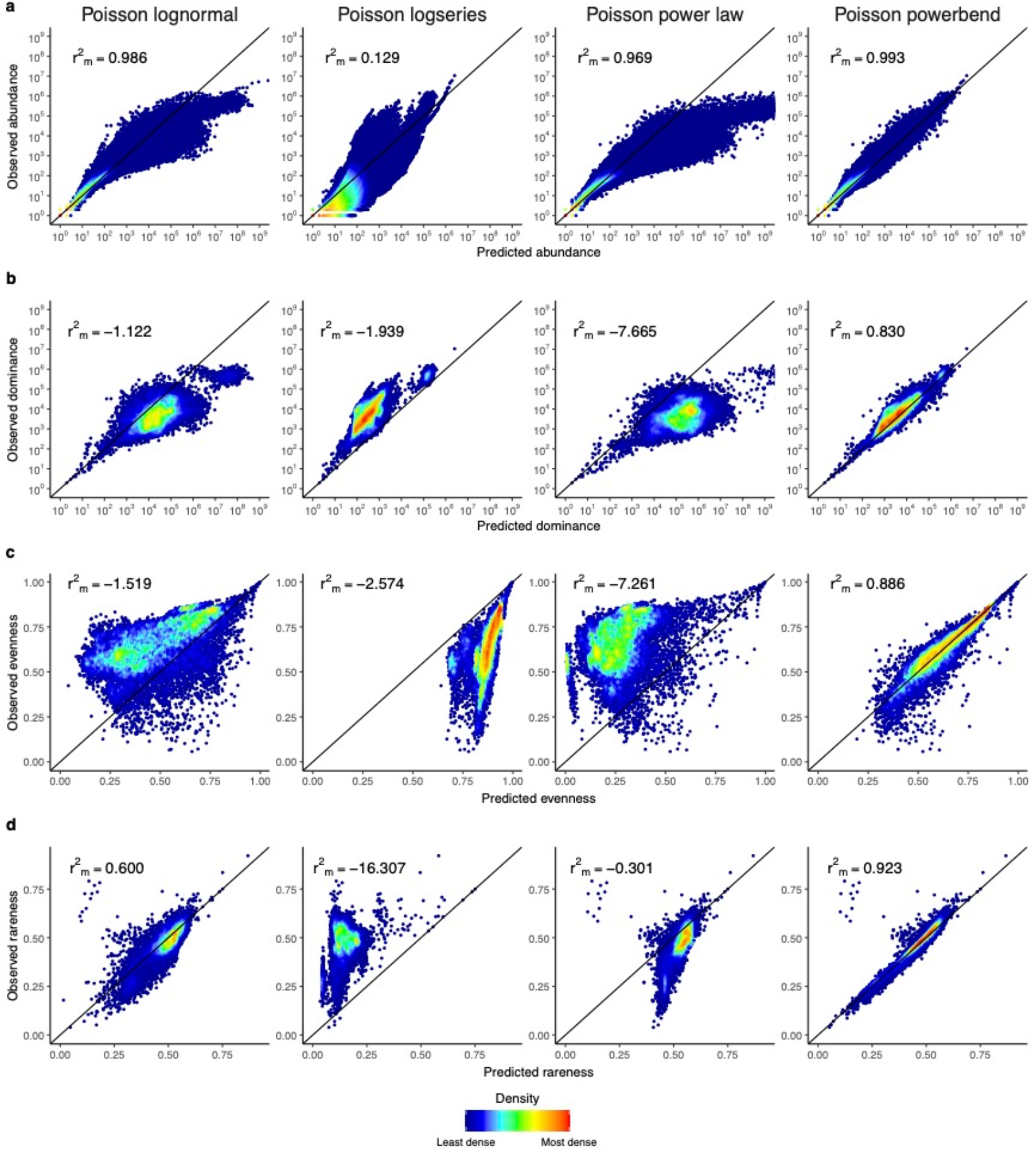
Goodness of fit of SAD models in 15,329 microbial communities. The layout of the figure is the same as that of Fig. 1. For illustrative purposes, the x-axis is truncated to 10^9^ in **a** and **b** because the power law overpredicts the abundance of the most abundant species.

We noted that Figure 2 of the study by Shoemaker et al. did not seem to show the overprediction of species abundance by the Poisson lognormal model ^3^, as we have demonstrated here. This discrepancy arises because the overprediction is most apparent in SADs with 16S rRNA reads exceeding 10^5^, and these SADs were excluded from Shoemaker et al.’s Figure 2 (Extended Data Fig. 2 and Supplementary Information).

### Powerbend accurately captures SAD skewness, evenness, and dominance while other models fail

SAD is often considered a weak test due to its limited ability to distinguish between ecological models using benchmarks such as AIC and R^2^ that measure the overall fit to the species abundance data^4^. One issue with the overall fit measurements is that they are heavily weighted by rare species that make up most of the data points. To overcome this problem, additional SAD metrics can be used to evaluate SAD models^3,20^. They encompass various features of the SAD including rareness (the asymmetry in the distribution of species abundance as a measure of rarity), evenness (the level of uniformity in species abundance) and dominance (N_max_, the abundance of the most abundant species). A good SAD model should not only capture the overall species abundance distribution in the raw data but also accurately reflect key scalar metrics such as evenness, rarity, and richness. Figs. 1b-d and Figs. 2b-d show that the powerbend model performs very well in capturing these additional SAD metrics of animal, plant, and microbial communities, while the other models all perform poorly. For example, although Poisson lognormal provides a good overall fit to the species abundance data in microbial communities (Fig. 2a), it fits poorly to the SAD dominance (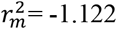, Fig. 2b), evenness (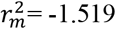, Fig. 2c) and rareness (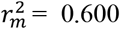, Fig. 2d). This result demonstrates again that the powerbend model is overwhelmingly superior to the other SAD models.

### Total number of Earth microbial species estimated using the lognormal SAD model can be orders of magnitude off

An exhaustive inventory of microbial communities remains impractical. Because of the parametric nature of SAD models, they have been applied to estimate the total number of species in a microbial community (including both observed and unobserved species). For example, lognormal has been used to estimate the total number of microbial species on Earth given the abundance of the most abundant species (N_max_) and the total number of individuals in the community (N) ^20,21^. However, we have shown that empirical microbial SADs deviate significantly from the lognormal model, especially for the most abundant species (Fig. 2b). Such model misspecification can lead to errors and biases in the estimated biodiversity.

To test this hypothesis, we simulated communities using the powerbend model, identified as the best model for microbial communities in this study, as the true SAD model. We found that the number of species estimated using lognormal varies greatly from the true value, and the direction and magnitude of the relative error are influenced by the *s* and λ parameters of the powerbend model (Fig. 3). For any given microbial community surveyed by 16S rRNA sequencing, the *s* and λ parameters of the powerbend model can be estimated, allowing for the estimation of species numbers in that community. However, no such survey exists for the global microbial community, and thus, the *s* and *λ* parameters for the global microbial community powerbend model remain unknown. Therefore, we systematically explored various combinations of these two parameters to yield the previously reported N_max_ and N values (N_max_=2.0×10^28^ and N=3.2×10^30^) for the Earth microbial community^20^(Table 2). Our findings revealed that N_max_ and N values alone do not suffice to ascertain the total number of microbial species. Notably, the global species number, highly contingent on the *s* parameter, can vary by up to 20 orders of magnitude. Therefore, we argue against estimating microbial species number via the lognormal model, as has been done previously^20,21^.

**Table 2.** Combinations of powerbend *s* and *λ* parameters that all produce previously reported N_max_ = 2.0×10^28^ and N = 3.2×10^30^ for the global Earth microbial community.

| $s$ | $\lambda$ | microbial species number |
| --- | --- | --- |
| 1.0 | $2.80 \times 10^{-28}$ | $5.60 \times 10^4$ |
| 1.05 | $2.70 \times 10^{-28}$ | $3.90 \times 10^5$ |
| 1.1 | $2.60 \times 10^{-28}$ | $4.80 \times 10^6$ |
| 1.15 | $2.60 \times 10^{-28}$ | $7.40 \times 10^7$ |
| 1.2 | $2.50 \times 10^{-28}$ | $1.30 \times 10^9$ |
| 1.25 | $2.40 \times 10^{-28}$ | $2.30 \times 10^{10}$ |
| 1.3 | $2.40 \times 10^{-28}$ | $4.50 \times 10^{11}$ |
| 1.35 | $2.30 \times 10^{-28}$ | $8.70 \times 10^{12}$ |
| 1.4 | $2.20 \times 10^{-28}$ | $1.70 \times 10^{14}$ |
| 1.45 | $2.10 \times 10^{-28}$ | $3.40 \times 10^{15}$ |
| 1.5 | $2.10 \times 10^{-28}$ | $6.80 \times 10^{16}$ |
| 1.55 | $2.00 \times 10^{-28}$ | $1.30 \times 10^{18}$ |
| 1.6 | $1.90 \times 10^{-28}$ | $2.70 \times 10^{19}$ |
| 1.65 | $1.80 \times 10^{-28}$ | $5.30 \times 10^{20}$ |
| 1.7 | $1.70 \times 10^{-28}$ | $1.00 \times 10^{22}$ |
| 1.75 | $1.60 \times 10^{-28}$ | $1.90 \times 10^{23}$ |
| 1.8 | $1.50 \times 10^{-28}$ | $3.60 \times 10^{24}$ |

**Fig 3.**
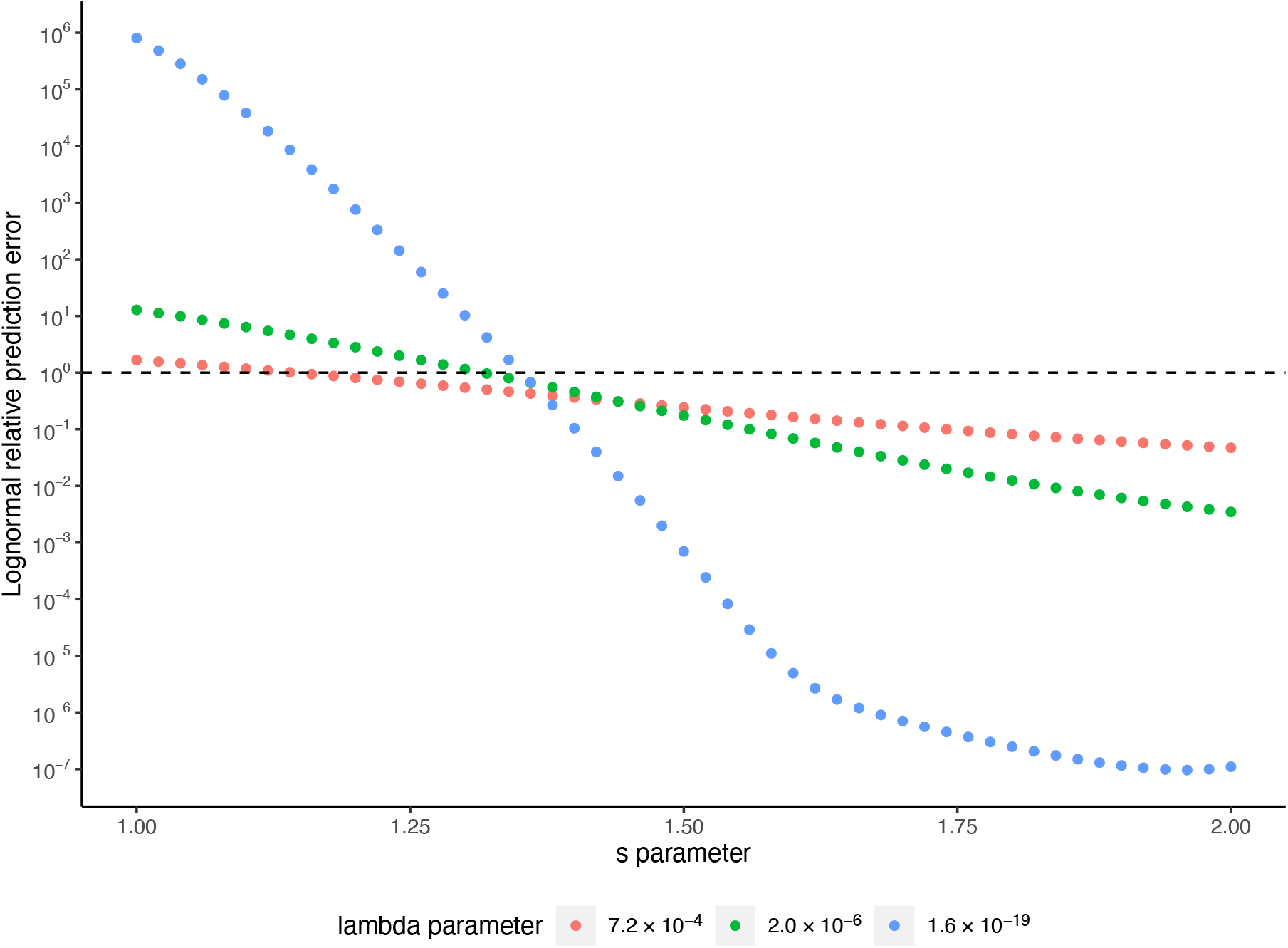
Microbial biodiversity estimated using lognormal can be orders of magnitude off. The number of species is predicted using the lognormal model and compared against the true number of species in the SAD simulated using powerbend models. The relative prediction error, defined as the predicted species number divided by the true species number, is plotted against the powerbend’s *s* parameter. Three different λ parameters were selected to represent the 2.5%, 50.0% and 97.5% percentiles of the λ parameters estimated from the 15,329 microbial communities.

## Discussion

The distribution of species abundance stands as one of the oldest, most universal, and fundamental laws in ecology. Despite decades of research and the development of numerous models, a universally accepted SAD model applicable across the tree of life remains elusive. This is partially due to the relatively low number of species within animal and plant communities limiting their power to distinguish between different SAD models. This challenge can be mitigated by testing SAD models within microbial communities, which typically exhibit significantly greater species richness. In this study, for instance, microbial communities have a median value of 3,246 species per SAD, facilitating more robust model testing. We boosted the power of SAD testing by concurrently evaluating models across animal, plant, and microbial communities, covering a range of abundance scales spanning 7 orders of magnitude. We also incorporated additional SAD metrics, including evenness, dominance, and rareness. Moreover, we explicitly modeled the sampling effort of surveys, an important but often overlooked factor in SAD model testing ^22^. As a result, we were able to reject commonly used lognormal, logseries, and power law models, establishing the powerbend as an emerging unifying SAD model. Our simulations show that when sampling or species number is sufficient, SAD data have enough power to distinguish SAD models and recover the true model (Extended Data Fig. 1). Conversely, poor sampling or the small number of species tends to favor simpler models such as the logseries and lognormal over the powerbend (Extended Data Fig. 1). Therefore, we concluded that the superiority of the powerbend model is unlikely to be a result of poor sampling. While the success of the powerbend model does not by itself prove the existence of universal ecological principles, it provides key evidence supporting this hypothesis.

It is important to point out that powerbend is in essence a “bended” power law^18^. While the abundance distribution of most species within a community follows a power law, the bending in the powerbend model imposes an upper limit on the abundance of dominant species. This bending reflects the inherent constraint on the population size of a community^15^ and is a key feature that distinguishes powerbend as a superior model to the power law. In contrast, the power law fits SADs poorly because it consistently overestimates the abundance of dominant species within a community (Figs. 1a and b, 2a and b).

The existence of a unifying SAD model suggests some common mechanism at work. Although a-mechanistic models such as the powerbend model do not explicitly incorporate the underlying mechanisms or processes, they can still be useful for discerning and revealing dominant mechanistic drivers of the ecosystem ^15,23^. For example, because the powerbend model was derived by modeling interspecific trait variation^16,17^, it supports the idea that species competition for resources, a deterministic mechanism, is important in driving the community assembly. Therefore, our result challenges purely neutral models such as Hubbell’s neutral theory, which predicts logseries SAD. On the other hand, the fact that the powerbend model is derived using a MaxEnt framework implies that stochasticity also plays an important role in shaping macroecological patterns.

The *s* parameter in the powerbend model can be conceptualized as representing the number of limiting resources driving the community assembly^23^. Our study indicates that in animal and plant communities, the weighted mean of the *s* parameter is 0.9, close to 1.0 when the powerbend model degenerates into the logseries model. In this scenario, the total available energy can be considered the limiting resource driving the macroecological patterns^23^. In contrast, the mean *s* parameter of microbial communities is 1.6. This suggests that microbe communities are constrained by a greater number of limiting resources, likely because microbial surveys typically span a much broader taxonomic range than animal and plant surveys. A greater number of limiting resources provides more specialized opportunities for rare species to survive, resulting in more uneven SADs or a more pronounced ‘rare biosphere’ in microbial communities. The fractional nature of the *s* parameter also implies that limiting resources are hierarchically partitioned among community members. This aligns with findings showing that cross-feeding between species in a microbial community is a key driver of community assembly ^24^.

The prediction of the powerbend SAD model is rooted in a maximum entropy-based framework. The success of the powerbend SAD model provides evidence that the assembly of both microorganism and macroorganism communities adheres to the same governing principles. This study should stimulate additional testing of maximum entropy-based theories as potential unifying frameworks for understanding other macroecological patterns, including the species-area relationship and the metabolic rate-abundance relationship^15,23,25^.

## Methods

### SAD Data

To test the universality of SAD models in animals and plants, the species abundance data were downloaded from Baldridge et al. study^2^, which include the Breeding Bird Survey (BBS)^26^, Alwyn H. Gentry’s Forest Transect Data Set (GENTRY)^27^, the Mammal Community Database (MCDB)^28^, the Forest Inventory and Analysis (FIA)^29^ and SAD data compiled from literature for an assortment of taxa^30^. To test microbial SADs, the bacterial and archaeal species abundance data were downloaded from Shoemaker et al. study^3^, which include data from the Earth Microbiome Project (EMP)^31^, the first phase of Human Microbiome Project (HMP1)^32^ and the MG-RAST repository (MGRAST)^33^. Our simulations showed that when the number of observed species in a SAD is small or when the sampling effort is low, model selection using AIC does not have sufficient power to distinguish SAD models, often favoring the simpler models even when the simpler model is incorrect (Extended Data Fig. 1). The more complex the model is, the more data points (i.e., species) are required to recover the true model. To strike a balance between the number of SADs we can analyze and the power of model selection, and following the practice of the previous studies^1,2^, we filtered out SADs with less than 10 species to test base SAD models for animal and plant SAD data, and SADs with less than 100 Operational Taxonomic Units (OTUs) to test compound SAD models for microbial SAD data. In addition, in the microbial SAD data, we identified 687 outlier SADs where the number of doubleton species exceeded the number of singleton species by more than 10-fold (Extended Data Fig. 3). These outliers all originated from the EMP dataset compiled by Shoemaker et al.^3^ but are no longer present in the current EMP dataset (year 2017 version). Consequently, we excluded these outliers from our analyses. This resulted in a total of 13,819 animal and plant SADs and 15,329 microbial SADs (see Extended Data Table 2).

The animal and plant dataset encompasses diverse taxonomic groups (see Extended Data Table 2). To ensure equitable representation across these groups, we applied weighting to each SAD based on the size of the dataset it originated from when assessing the frequency of a model being the best by AIC or its goodness of fit. For instance, a SAD within the Mammal Community Database (MCDB, 103 samples) carried 26.9 times the weight of a SAD from the Breeding Bird Survey (BBS, 2,769 samples).

### SAD models

The lognormal SAD model has the probability density function:

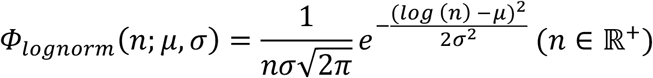

where *n* represents the species abundance, and *μ* and *σ* are the mean and standard deviation of log-transformed *n*, respectively. The logarithm is calculated with the natural base *e*. Given that the lognormal distribution is continuous while the SAD is inherently discrete, it is necessary to convolute it with a sampling error when fitting it to SAD data.

The logseries distribution has the probability mass function:

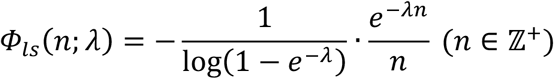

where *λ* is the exponential rate at which the numerator decays.

The power law distribution has the probability mass function:

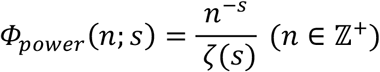

where *s* > 1 is the scaling parameter that controls the distribution’s decay rate, and *ζ*(*s*) is the Riemann zeta function, serving as a normalization factor to ensure the sum of the probabilities over all possible values of *n* equals 1.

The powerbend distribution takes a more general form compared to the logseries and power law distributions. It is a hybrid of a power law and an exponential function, in which the exponential function bends the power law by setting an upper bound to the power law^18^. It has the probability mass function:

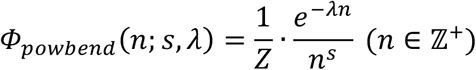

where *s* is the order of the denominator, λ is the exponential rate at which the numerator decays, and *Z* is the probability normalizer. It should be noted that the powerbend distribution is a generalized distribution that can degenerate into the logseries, geometric, broken-stick, and power law distributions by setting its parameters to certain fixed values (Supplementary Information).

### Modeling sampling effort in 16S rRNA survey

To relate the number of observed reads in 16S rRNA profiling data to the number of individual cells in the community, we explicitly modeled the sampling effort by convoluting a sampling error to the base SAD models. The 16S rRNA profile of a community typically comprises thousands of reads. Assuming no bias, the sampling error can be approximated by a Poisson distribution whose mean represents the expected number of reads given the number of individual cells. We used parameter *η* to denote the expected number of reads per individual cell, which represents the sampling effort. By convoluting the sampling error to the base SAD models, we derived the sampling-explicit SAD models defined on non-negative integers *k* ∈ 0 ∪ ℤ^+.^:

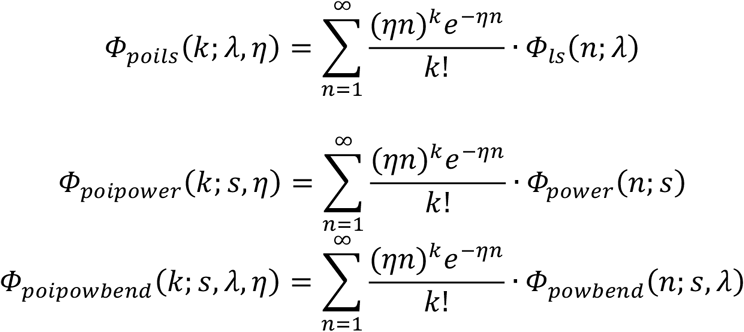

It should be noted that for lognormal distribution, because the number of individuals *n* and the sampling effort *η* always appear together in the formula as their product, there is no way to disentangle the two parameters. Therefore, the sampling effort *η* is fixed to 1 and the probability mass function of the Poisson lognormal distribution has the same number of parameters as the base distribution:

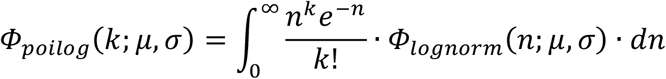

In practice, there may be bias in 16S rRNA sequencing resulting from biases in DNA extraction and PCR amplification. Thus, a Poisson sampling error may not be sufficient to accurately relate the sequence reads to the number of individual cells. Because the direction and magnitude of such bias are often unknown, its effect on the distribution of sequence reads can be seen as an inflation of the variance or over-dispersion without changing the mean. As a result, we used the negative binomial distribution to model such bias^34^, and derived the corresponding probability mass function for SAD models:

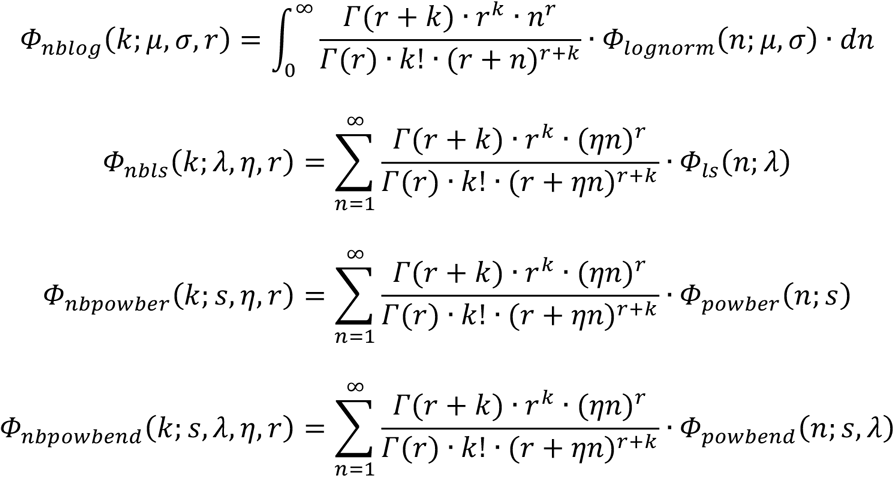

In the above equations, *r* measures the degree of over-dispersion or bias in the negative binomial distribution and *Γ* is the gamma function.

Using 200 randomly selected microbial SADs, we compared the use of Poisson and negative binomial distributions to model the sampling effort in the 16S rRNA sequencing pipeline. Our result indicated that SAD models with the Poisson error structure are superior to those with the negative binomial error structure (Extended Data Table 6). Therefore, we used the Poisson error structure for testing the full microbial SAD dataset.

Extended Data Table 1 lists the models tested in this study and their free parameters.

### Fitting SAD models to empirical data

Because all SAD models (both base models and their sampling-explicit counterparts) are formulated as probability distributions, we fitted them to the observed abundances in empirical data using the maximum likelihood (ML) framework. Because species with zero observations are not recorded in empirical data, the likelihood of a single species with *k* observations (denoted as *L*(*k*)) for sampling-explicit SAD models is normalized by the cumulative probability of any positive observation:

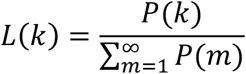

In the above formula, *P*(*k*) is the probability mass function for one of the sampling-explicit SAD models described in the previous section. Following the practice of ^1–3^, we compared the performance of SAD models using AIC, a commonly used metric in SAD model testing.

### Evaluating the goodness of fit of SAD models

The goodness of fit of SAD models was evaluated by comparing predicted and observed species abundances in the rank-abundance distribution (RAD). Briefly, to generate the expected RAD for a SAD with *S* observed species, we placed *S* quantiles on the cumulative distribution function (CDF) of the fitted SAD model, evenly dividing the cumulative probability (the y-axis of the CDF curve). For comparison between the expected and the observed RADs, we used the modified coefficient of determination around the 1:1 line 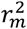 as described in previous studies^1,3,19^ to quantify the goodness of fit:

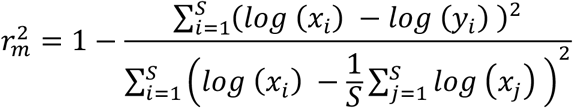

In the above formula, *x* and *y* are the observed and the predicted abundances (e.g., number of reads in 16S rRNA profiling data), respectively. The subscripts denote the rank of species, and *S* is the total number of observed species in the SAD. Because the coefficient of determination here is calculated against the 1:1 line with a fixed intercept of 0, it is possible that its value drops below zero, which indicates a poor fit.

The value of 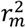 reaches 1 if and only if the observed and the predicted abundances match perfectly, indicating a perfect fit of SAD. To determine if the 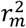 of a model fitted to a SAD (observed 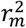) is statistically significantly lower than 1, we established a baseline distribution of 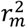 values from 1,000 iterations of Monte-Carlo simulation. In each iteration, we simulated a RAD based on the fitted SAD model and calculated 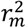 between the simulated RAD and the expected RAD. We then employed a one-sided test to assess the lack-of-fit by calculating the empirical frequency at which the baseline 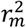 values were smaller than the observed 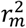.

In addition, the goodness of fit of SAD models was evaluated by comparing observed and predicted SAD evenness, rareness, and dominance. Evenness of a SAD was measured using Shannon’s evenness E_H_, also known as Pielou’s evenness J^35^. Rareness was measured by the log-modulo of skewness of a SAD, as described in^3,20^, with log-transformed species abundances utilized for the skewness calculation. Dominance is measured simply as the abundance of the most abundant species (N_max_) in the SAD, also as described in^3,20^.

### Evaluating the effects of observed species number and sampling effort on SAD model selection

We investigated the influence of the number of observed species on the efficacy of model selection by AIC. We generated simulated communities employing either a logseries (λ = 10^-2^) or powerbend (*s* = 1.5, λ = 10^-2^) SAD model. The simulated communities varied in the observed species number, ranging from 10 to 100 species per community. For each level of species count, we simulated 100 communities under each SAD model. Subsequently, we fitted the logseries and powerbend models to the simulated data and recorded the frequency at which each model emerged as the best-fit model by AIC.

Likewise, we examined the effect of sampling effort on SAD model selection. We conducted simulations using communities generated according to either a powerbend (*s* = 1.5 and λ = 10^-5^) or lognormal (*μ* = 0 and *σ* = 3.25) SAD model. For each SAD model, we simulated 100 communities, each consisting of 10^4^ species, by randomly drawing species abundances from the respective SAD distribution. Next, we simulated the sampling process by randomly drawing the number of observations for each species from a Poisson distribution, whose mean was set to be the product of the species abundance and the sampling effort. We simulated 9 levels of sampling effort, evenly spaced from 10^-1^ to 10^-5^ on a log-scale. Subsequently, we fitted the Poisson powerbend and Poisson lognormal models to the simulated data and recorded the frequency at which each model appeared as the best model at each level of sampling effort.

### Evaluating the effect of OTU sequence identity threshold on SAD model selection

To investigate the effect of OTU sequence identity threshold on SAD model selection, we clustered OTU at different identity thresholds for the 565 SADs from the HMP1 dataset using Mothur (version 1.48)^36^. Specifically, we extracted all aligned 16S rRNA gene sequences in the HMP1 dataset from the summary table of the Mothur pipeline. We calculated the pairwise distance between unique sequences using the function “dist.seqs” (with arguments cutoff=0.20 and output=lt) and clustered the unique sequences with the average neighbor algorithm using the function “cluster” (with arguments: method=average and cutoff=0.20). OTU tables were generated at the sequence identity threshold of 95% and 99% using the function “make.shared” (with argument: label=0.01-0.05). We fitted SAD models to these two datasets, conducted model selection through AIC, and compared the outcome with that of the original HMP1 dataset.

### Evaluating the effect of 16S rRNA gene copy number (GCN) correction on SAD model selection

Because 16S rRNA GCN variation can bias the species composition estimated using 16S rRNA read counts^37^, we assessed its impact on the outcome of SAD model selection by performing model selection on the 565 SADs from the HMP1 dataset. We predicted GCN for each OTU and then fitted models on the species abundance data corrected for GCN variation. Specifically, we selected the most abundant sequence in each OTU as its representative sequence. We then predicted the 16S GCN of each OTU using *RasperGade16S*^38^, an R package that predicts 16S GCN based on phylogenetic relatedness. In SAD model fitting, we modeled 16S GCN as an OTU-specific multiplier to the sampling effort *η*. Consequently, only SAD models with a Poisson sampling error structure were included in this analysis. Model selection was performed through AIC, and the outcomes were compared with those obtained without 16S GCN correction.

### Evaluating the effect of SAD model misspecification on biodiversity estimation

To study the impact of model misspecification on biodiversity estimation using the lognormal model, we simulated communities whose SAD follows the powerbend model. 95% of the *s* and *λ* parameters estimated from the 15,329 microbial communities were within the range of 1.2-1.8 (median: 1.6) and 7.2×10^-4^-1.6×10^-19^ (median: 2.0×10^-6^), respectively. Therefore, we simulated SADs with *s* parameters ranging from 1.0 to 2.0 at an interval of 0.02 and the λ parameters fixed to 7.2×10^-4^, 2.0×10^-6^ or 1.6×10^-19^. We predicted the species number based on the abundance of the most abundant species in the community (N_max_) and the total number of individuals (N), using the lognormal model following the method of Curtis et al.^21^. We calculated the relative prediction error as the ratio of the predicted species number to the true species number used in our simulations. To find possible *s* and λ parameters that yield communities aligning with the published estimates of N_max_ (2.0×10^28^) and N (3.2×10^30^) for the Earth microbial community^20^, we first fixed the *s* parameter to a specific value (e.g., 1.2) and then optimized the λ parameter to minimize the sum of squared differences between simulated and published estimates of N_max_ and N. This process was repeated across a range of *s* parameter values, allowing us to identify feasible combinations of *s* and *λ* parameters consistent with the published estimates of N_max_ and N for the Earth microbial community, and the corresponding species numbers.

## Data and code availability

The SAD data that support the findings of this study, and code and instructions (a Readme file) for replicating the analyses are available at figshare: doi:10.6084/m9.figshare.25711257. The R package *microSAD* can be downloaded from Github: https://github.com/wu-lab-uva/microSAD.

## Acknowledgements

We thank Research Computing at The University of Virginia for providing computational resources and technical support that have contributed to the results reported within this publication.

## Author contributions

YG and AA developed the R package *microSAD*. YG, AA and MW conducted data analyses. YG and MW wrote the manuscript. All authors read and approved the final manuscript.

## Ethics declarations

Competing interests

The authors declare no competing interests.

Corresponding author

Correspondence to Martin Wu

## References

1. White, E. P., Thibault, K. M. & Xiao, X. Characterizing species abundance distributions across taxa and ecosystems using a simple maximum entropy model. Ecology 93, 1772–1778 (2012).

2. Baldridge, E., Harris, D. J., Xiao, X. & White, E. P. An extensive comparison of species-abundance distribution models. Peerj 4, e2823 (2016).

3. Shoemaker, W. R., Locey, K. J. & Lennon, J. T. A macroecological theory of microbial biodiversity. Nat Ecol Evol 1, 0107 (2017).

4. McGill, B. J. et al. Species abundance distributions: moving beyond single prediction theories to integration within an ecological framework. Ecol Lett 10, 995–1015 (2007).

5. Sogin, M. L. et al. Microbial diversity in the deep sea and the underexplored “rare biosphere.” Proc. Natl Acad. Sci. 103, 12115–12120 (2006).

6. Fisher, R. A., Corbet, A. S. & Williams, C. B. The relation between the number of species and the number of individuals in a random sample of an animal population. J Animal Ecol 12, 42 (1943).

7. Preston, F. W. The commonness, and rarity, of species. Ecology 29, 254–283 (1948).

8. MacArthur, R. H. On the relative abundance of bird species. Proc. Natl Acad. Sci. 43, 293–295 (1957).

9. Motomura, I. A statistical treatment of ecological communities. Zoological Magazine 44, 379–383 (1932).

10. Frontier, S. Diversity and structure in aquatic ecosystems. Oceanography and Marine Biology 23, 253–312 (1985).

11. Hubbell, S. P. The Unified Neutral Theory of Biodiversity and Biogeography. (Princeton University Press, Princeton, 2001).

12. Volkov, I., Banavar, J. R., Hubbell, S. P. & Maritan, A. Neutral theory and relative species abundance in ecology. Nature 424, 1035–1037 (2003).

13. Alroy, J. The shape of terrestrial abundance distributions. Sci Adv 1, e1500082 (2015).

14. Leigh, E. G. Tropical Forestecology A View from Barro Colorado Island. (Oxford University Press, Oxford, 1999).

15. Harte, J., Zillio, T., Conlisk, E. & Smith, A. B. Maximum entropy and the state-variable approach to macroecology. Ecology 89, 2700–2711 (2008).

16. Pueyo, S., He, F. & Zillio, T. The maximum entropy formalism and the idiosyncratic theory of biodiversity. Ecol. Lett. 10, 1017–1028 (2007).

17. Bertram, J., Newman, E. A. & Dewar, R. C. Comparison of two maximum entropy models highlights the metabolic structure of metacommunities as a key determinant of local community assembly. Ecol Model 407, 108720 (2019).

18. Pueyo, S. Diversity: between neutrality and structure. Oikos 112, 392–405 (2006).

19. Xiao, X., McGlinn, D. J. & White, E. P. A strong test of the maximum entropy theory of ecology. Am Nat 185, E70–E80 (2015).

20. Locey, K. J. & Lennon, J. T. Scaling laws predict global microbial diversity. Proc. Natl Acad. Sci. 113, 5970–5975 (2016).

21. Curtis, T. P., Sloan, W. T. & Scannell, J. W. Estimating prokaryotic diversity and its limits. Proc. Natl Acad. Sci. 99, 10494–10499 (2002).

22. McGill, B. J. et al. Species abundance distributions: moving beyond single prediction theories to integration within an ecological framework. Ecol Lett 10, 995–1015 (2007).

23. Harte, J. & Newman, E. A. Maximum information entropy: a foundation for ecological theory. Trends Ecol Evol 29, 384–389 (2014).

24. Goldford, J. E. et al. Emergent simplicity in microbial community assembly. Science 361, 469–474 (2018).

25. Shade, A. et al. Macroecology to unite all Life, large and small. Trends Ecol Evol 33, 731–744 (2018).

26. Ziolkowski, D., Lutmerding, M., Aponte, V. I. & Hudson, M.-A. R. North American Breeding Bird Survey Dataset (1966-2021). doi:10.5066/P97WAZE5.

27. Phillips, O. & Miller, J. S. Global Patterns of Plant Diversity: Alwyn H. Gentry Forest Transect Data Set. (Missouri Botanical Garden Press, St. Louis, 2002).

28. Thibault, K. M., Supp, S. R., Giffin, M., White, E. P. & Ernest, S. K. M. Species composition and abundance of mammalian communities. Ecology 92, 2316–2316 (2011).

29. Forest inventory and analysis national core field guide (Phase 2 and 3). https://www.fs.usda.gov/research/programs/fia.

30. Baldridge, E. Community abundance data. https://figshare.com/articles/dataset/Community_abundance_data/769251/1.

31. Gilbert, J. A., Jansson, J. K. & Knight, R. The Earth Microbiome project: successes and aspirations. Bmc Biol 12, 69 (2014).

32. Turnbaugh, P. J. et al. The human microbiome project. Nature 449, 804–810 (2007).

33. Meyer, F. et al. The metagenomics RAST server – a public resource for the automatic phylogenetic and functional analysis of metagenomes. Bmc Bioinformatics 9, 386 (2008).

34. Bliss, C. I. & Fisher, R. A. Fitting the negative binomial distribution to biological data. Biometrics 9, 176 (1953).

35. Pielou, E. C. Ecological Diversity. (John Wiley & Sons, New York, 1975).

36. Schloss, P. D. et al. Introducing mothur: open-source, platform-independent, community-supported software for describing and comparing microbial communities. Appl. Environ. Microbiol. 75, 7537–41 (2009).

37. Kembel, S. W., Wu, M., Eisen, J. A. & Green, J. L. Incorporating 16S gene copy number information improves estimates of microbial diversity and abundance. PLoS computational biology 8, e1002743 (2012).

38. Gao, Y. & Wu, M. Accounting for 16S rRNA copy number prediction uncertainty and its implications in bacterial diversity analyses. ISME Commun. 3, 59 (2023).

