## Supplementary Information for "A unifying model of species abundance distribution"

**Supplementary Methods**

Replicating lognormal predictions in Figure 2 of the Shoemaker et al. 2017 study

To replicate Figure 2 in the Shoemaker et al. (2017) study, we downloaded and ran the Python scripts from the original study (https://github.com/LennonLab/MicrobialBiodiversityTheory). During this process, we identified a problem in their study.

To evaluate the goodness of fit, Shoemaker et al. fitted over 20,000 SADs using different models to predict species abundances, and then compared the observed versus predicted abundances. However, the Python script set a 21-second time limit for predicting species abundance for each community. Predictions using the lognormal and Zipf power law models were more time-consuming than those using the logseries and broken-stick models. As a result, only 26% of SADs had successful species abundance predictions for either the lognormal or Zipf model, compared to 47% and 89% for the broken-stick and logseries models, respectively. This meant that different SAD models were evaluated using different sets of SADs, resulted in unfair comparisons. SADs with species abundances exceeding $10^5$ were excluded from the lognormal and Zipf models, because predicting species abundance for these SADs exceeded the 21-second time limit. This is evident in Figure 2 of the Shoemaker et al. study, where the lognormal and Zipf models were evaluated with SAD data with a maximum observed species abundance of $10^5$, while the logseries and broken-stick models were evaluated with SAD data with a maximum observed species abundance of $10^6$.

In our study, the overprediction by the lognormal model is most apparent when species abundance exceeds $10^5$ (Fig. **2a**). Therefore, we suspected the lognormal predictions presented in Figure 2 of the Shoemaker et al. study were misleading because they had excluded the overpredicted species. To illustrate our point, we randomly selected 600 SADs (200 each from the HMP1, EMP, and MG-RAST datasets) and predicted species abundances using Shoemaker et al.'s Python scripts. We ran the scripts using either the default 21-second time limit (as in the original Shoemaker et al. study) or an extended time limit of 3 hours. We then plotted the observed species abundances against the predicted species abundances under both conditions, as well as the predictions made using the R package *microSAD* for the same 600 SADs. As shown in Extended Data Fig. 2, the overprediction by the lognormal model becomes apparent when more SADs are included in the figure.

Derivation of various SAD models from the powerbend model

The powerbend distribution is a hybrid of a power law and an exponential function, in which the exponential function bends the power law by setting an upper bound to the power law [1]. It has the probability mass function:

$$\Phi_{powbend}(n; s, \lambda) = \frac{1}{Z} \cdot \frac{e^{-\lambda n}}{n^s} \ (n \in \mathbb{Z}^+)$$

where $s$ is the order of the denominator, λ is the exponential rate at which the numerator decays, and $Z$ is the probability normalizer that is equal to polylogarithm $Li_s(e^{-\lambda})$.

Logseries

Logseries was first introduced by Fisher [2]. Under Fisher's parameterization, the logseries has the probability mass function:

$$\Phi_{ls}(n; \alpha, x) = \alpha \frac{x^n}{n}$$

where α and $x$ are constants derived from the total number of species and individuals.

Under an alternative parameterization, the logseries distribution has the probability mass function {Harte, 2008}:

$$\Phi_{ls}(n; \lambda) = -\frac{1}{\log(1 - e^{-\lambda})} \cdot \frac{e^{-\lambda n}}{n} \ (n \in \mathbb{Z}^+)$$

It is easily shown that when the powerbend $s$ parameter is fixed to 1, powerbend becomes logseries.

Power law

Power law distributions are a set of distributions that are commonly used to describe highly uneven distributions. In terms of species abundance, the power law distribution has the probability mass function:

$$\Phi_{power}(n; s) = \frac{n^{-s}}{\zeta(s)} \ (n \in \mathbb{Z}^+)$$

where $s > 1$ is the shape parameter that controls the distribution's decay rate, and $\zeta(s)$ is the Riemann zeta function, serving as a normalization factor to ensure the sum of the probabilities over all possible values of $n$ equals 1.

It is also easily shown that when the powerbend λ parameter is fixed to 0, powerbend becomes power law.

Broken stick

Originally developed by MacArthur [3], the Broken stick SAD is a continuous SAD and has the probability density function [4]:

$$\Phi_{broken-stick}(n;\ S, N) = \frac{S-1}{N}\left(1-\frac{n}{N}\right)^{S-2} \quad (n \in \{0, \mathrm{N}\})$$

where $S$ and $N$ are the total number of species and individuals in the sampled community, respectively. For discrete abundances, one can start from the asymptotic relation when $S$ and $N$ are both large[4]:

$$P(x \geq n) = e^{-\frac{S}{N}n}\ (n \in \{0\} \cup \mathbb{Z}^{+})$$

The probability mass function can then be derived as one for geometric distribution:

$$P(x = n) = P(x \geq n) - P(x \geq n+1) = \left(1 - e^{-\frac{S}{N}}\right)e^{-\frac{S}{N}n}\ (n \in \{0\} \cup \mathbb{Z}^{+})$$

Because in practice a species with an abundance of 0 will not be observed in the data, the probability mass function of the Broken stick model will be normalized to:

$$\Phi_{broken-stick}(n;\ \lambda) = \frac{1}{e^{\lambda}-1}e^{-\lambda n}\ (n \in \mathbb{Z}^{+})$$

where the exponential coefficient $\lambda = \frac{S}{N}$ is fixed.

When the powerbend $s$ parameter is fixed to 0, powerbend becomes the geometric distribution, which well approximates the broken stick model.

Geometric series

In the ecological literature, the geometric series was probably the first SAD ever proposed[5]. The name may be confused with the geometric distribution, but the geometric series directly describes the rank abundance distribution (RAD), instead of SAD. It has been shown that for continuous abundances, the geometric series RAD leads to a truncated continuous power law SAD[4]. Following the same strategy, we can derive the probability mass function for discrete abundances. Briefly, the geometric series RAD depicts that the expected abundance of a species with rank $i$ decreases at a constant proportion $0<k<1$ with regard to the species with rank $i$-1:

$$\frac{n_i}{n_{i-1}} = k \ (i > 1)$$

Given that a sampled community has $S$ species and $N$ individuals, we have the constraint that

$$N = n_1 \sum_{i=1}^{S} k^{i-1}$$

The constraint yields that the expected abundance of a species with rank $i$ is:

$$n_i = \frac{1-k}{1-k^S} k^{i-1} N$$

In the case when $S$ and $N$ are large, the survival function (upper tail cumulative distribution function) of a large species abundance $n$ can be written as:

$$P(x \geq n) = \frac{j}{S} \ (n \gg 1)$$

where $j$ satisfies the equation:

$$n_j = \frac{1-k}{1-k^S} k^{j-1} N = n$$

Solving the equation yields the survival function as:

$$P(x \geq n) = \frac{\log(k) + \log(n) + \log\left(\frac{1-k^S}{N(1-k)}\right)}{S \log(k)}$$

Then the probability mass function of the species abundance can be derived as:

$$P(x = n) = P(x \geq n) - P(x \geq n + 1) = \frac{\log\left(\frac{n}{n+1}\right)}{S \log(k)} = -\frac{\log\left(1 + \frac{1}{n}\right)}{S \log(k)}$$

Because we assume that $n \gg 1$, the probability mass function of the species abundance can be approximated by:

$$P(x = n) = -\frac{n^{-1}}{S \log(k)}$$

As a result, a geometric series RAD would roughly corresponds to a truncated power law SAD that has the probability mass function:

$$\Phi_{geomseries}(n) = \frac{1}{H_N} \cdot n^{-1} \ (n \in \{1,2,3, \dots, \mathrm{N}\})$$

where $H_N$ is the harmonic number that serves as the normalizing coefficient for the probability mass function. When the powerbend *s* parameter is fixed to 1, the λ parameter is fixed to 0, and the abundance has an upper limit *N*, powerbend describes the SAD that corresponds to the geometric series RAD.

### Extended data figures and tables

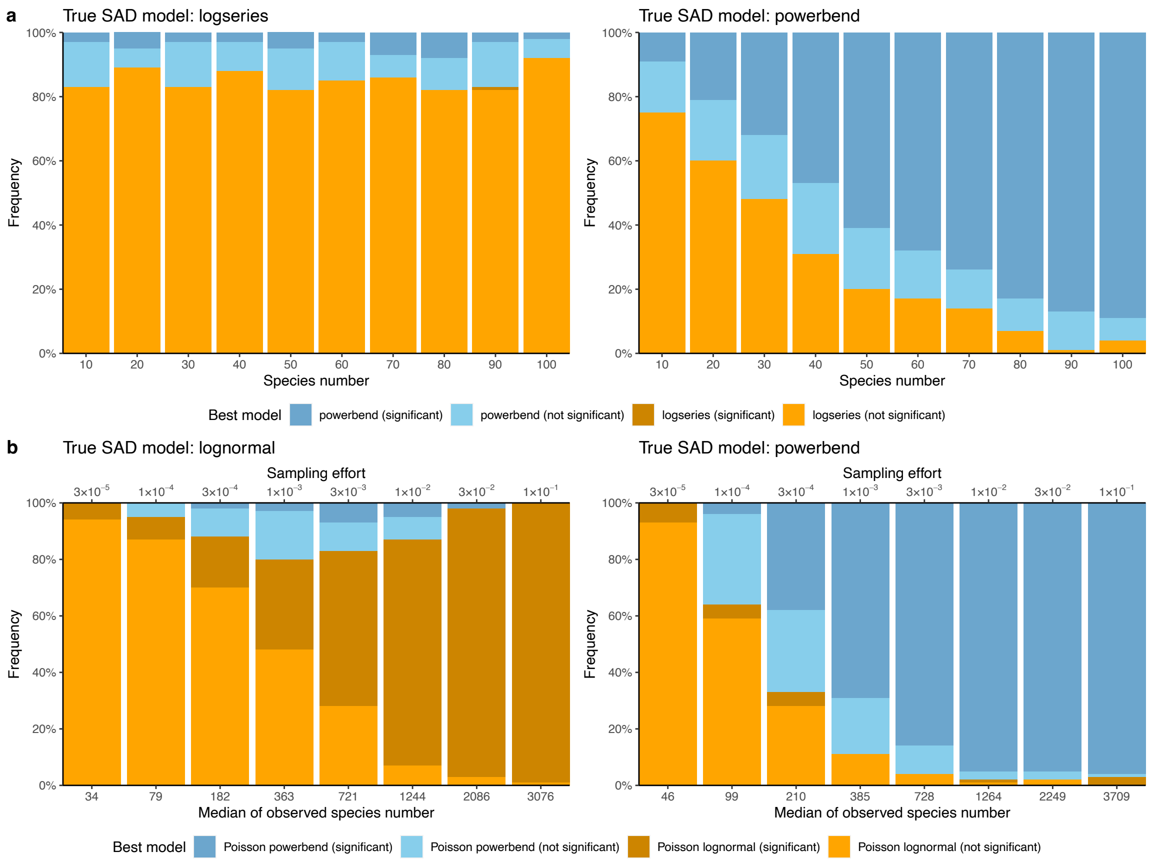


**Extended Data Figure 1 | Small observed species number and sampling effort favor simpler SAD models in model selection by AIC. a**, The frequency of the best model selected by AIC (y-axis) is plotted against the number of observed species in the SAD (x-axis), when the true SAD model is logseries (left) and powerbend (right). **b**, The frequency of the best model selected by AIC (y-axis) is plotted against different sampling efforts (x-axis), when the true SAD model is lognormal (left) and powerbend (right). The logseries, powerbend, Poisson lognormal and Poisson powerbend have 1, 2, 2 and 3 free parameters respectively. Therefore, the logseries model is simpler compared to powerbend, and the Poisson lognormal model is simpler compared to Poisson powerbend. A model is considered significantly superior if its AIC value is at least 2 points lower than that of the competing model.

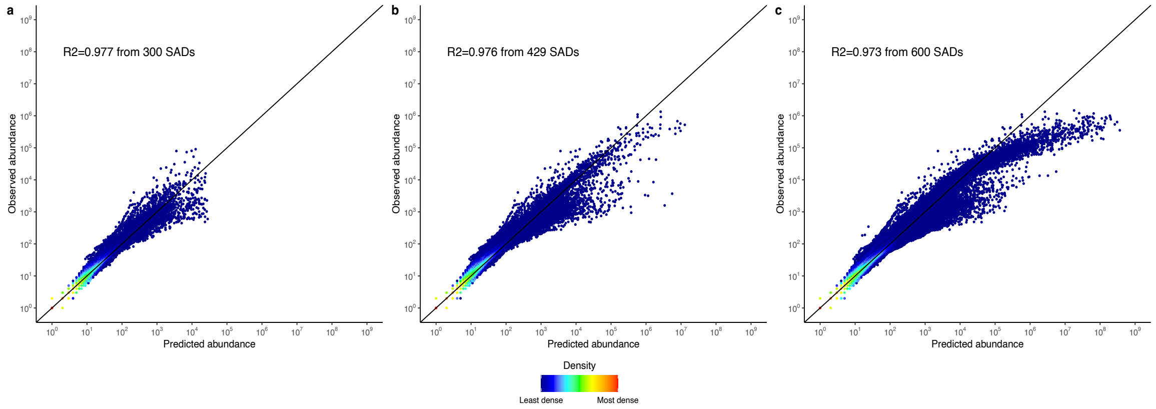


**Extended Data Figure 2 | Overprediction of the species abundance by the Poisson lognormal model.** The rational and methods used to create this figure are described in the Supplementary Information. Briefly, Shoemaker et al.'s Python code was used to fit 600 randomly sampled SADs (200 each from the HMP1, EMP and MG-RAST dataset) using the lognormal model to predict species abundances. The observed species abundances (y-axis) were plotted against the predicted species abundances (x-axis). Panel **a** shows 300 SADs for which the job of species abundance prediction took less than the 21-second time limit, as in the original Shoemaker et al. study. Panel **b** shows 429 SADs for which the job of species abundance prediction took less than 3 hours. Panel **c** shows all 600 SADs predicted in this study using the R package *microSAD*. Each data point represents a species. Note that overprediction is apparent when the observed species abundance exceeds $10^5$ (**b & c**), but these points are excluded from **(a)** due to the shorter time limit set in the original study. The color gradient represents dot density, with red indicating the highest density and dark blue the lowest. The black diagonal line represents a perfect 1:1 fit. Goodness of fit is determined by the modified coefficient of determination ($r_m^2$) against the 1:1 line, with the mean value shown.

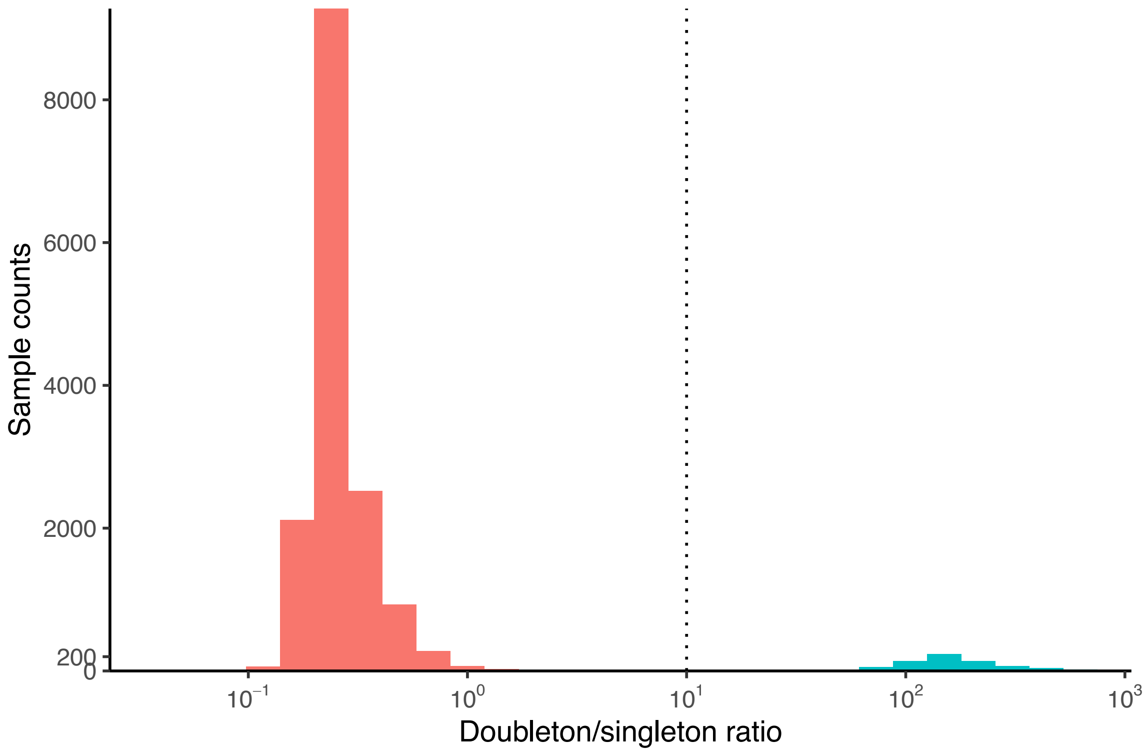


**Extended Data Figure 3 | Histogram of the doubleton/singleton ratio of 16,016 microbial SADs in the Shoemaker et al. study.** The doubleton/singleton ratio is calculated as the number of doubleton species divided by the number of singleton species in a SAD. Notably, there is a large gap separating the 687 SADs (highlighted in cyan) from the remainder of the 16,016 SADs. A doubleton/singleton ratio of 10 (indicated by the dotted line) was consequently employed to identify the outliers.

### Extended Data Table 1 | SAD models tested in this study

| SAD model | Sampling error structure | Free parameters |
|---|---|---|
| Lognormal | Poisson | 2 ($\mu$, $\sigma$) |
| | Negative binomial | 3 ($\mu$, $\sigma$, r) |
| Logseries | None | 1 ($\lambda$) |
| | Poisson | 2 ($\lambda$, $\eta$) |
| | Negative binomial | 3 ($\lambda$, $\eta$, r) |
| Power law | None | 1 ($s$) |
| | Poisson | 2 ($s$, $\eta$) |
| | Negative binomial | 3 ($s$, $\eta$, r) |
| Powerbend | None | 2 ($s$, $\lambda$) |
| | Poisson | 3 ($s$, $\lambda$, $\eta$) |
| | Negative binomial | 4 ($s$, $\lambda$, $\eta$, r) |

### Extended Data Table 2 | SAD datasets analyzed in this study

| | Dataset | Number of samples | Total |
|---|---|---|---|
| Animal and plant | Breeding Bird Survey (BBS) | 2,769 | 13,819 |
| | Alwyn H. Gentry's Forest Transect Data Set (GENTRY) | 220 | |
| | Mammal Community Database (MCDB) | 103 | |
| | Forest Inventory and Analysis (FIA) | 10,355 | |
| | Various taxa (reptiles etc) | 372 | |
| Bacteria and archaea | Earth Microbiome Project (EMP) | 13,772 | 15,329 |
| | Human Microbiome Project (HMP1) | 565 | |
| | MG-RAST repository (MGRAST) | 992 | |
| Total | | | 29,148 |

**Extended Data Table 3 | The average goodness of fit ($r_m^2$) for species abundance of the 13,819 animal and plant SADs**

| Model | Average $r_m^2$ (unweighted)* | $r_m^2$ values not statistically significant from 1.0 |
|---|---|---|
| Lognormal | 0.927 | 100% |
| Logseries | 0.822 | 95.3% |
| Power Law | 0.337 | 80.1% |
| Powerbend | 0.914 | 99.8% |

* Samples were not weighted by the size of datasets they originated from.

### Extended Data Table 4 | Frequencies of each SAD model being selected as the best model by AIC at different OTU thresholds

| OUT threshold (%) | Model | Sampling error structure | Best model | Statistically significant |
|---|---|---|---|---|
| 95 | Lognormal | Poisson | 20% | 8.10% |
| 95 | Logseries | Poisson | 8.30% | 0% |
| 95 | Power Law | Poisson | 6.40% | 0.20% |
| 95 | Powerbend | Poisson | 65.30% | 41.20% |
| 97 | Lognormal | Poisson | 18.20% | 6.20% |
| 97 | Logseries | Poisson | 3.50% | 0% |
| 97 | Power Law | Poisson | 7.60% | 0.20% |
| 97 | Powerbend | Poisson | 70.60% | 48.70% |
| 99 | Lognormal | Poisson | 14.70% | 5% |
| 99 | Logseries | Poisson | 0% | 0% |
| 99 | Power Law | Poisson | 10.10% | 0% |
| 99 | Powerbend | Poisson | 75.20% | 49.60% |

A best model is considered statistically significant when its AIC difference to the second-best model is greater than 2.

### Extended Data Table 5 | Frequencies of each SAD model being selected as the best model by AIC with and without 16S rRNA gene copy number (GCN) correction

| Model | Sampling error structure | Best model without GCN correction (statistically significant) | Best model with GCN correction (statistically significant) |
|---|---|---|---|
| Lognormal | Poisson | 18.23% (6.19%) | 17.52% (7.08%) |
| Logseries | Poisson | 3.54% (0%) | 5.49% (0%) |
| Power law | Poisson | 7.61% (0.18%) | 3.72% (1.06%) |
| Powerbend | Poisson | 70.62% (48.67%) | 73.27% (52.04%) |

A best model is considered statistically significant when its AIC difference to the second-best model is greater than 2.

**Extended Data Table 6 | Frequencies of SAD models with different sampling error structures being selected as the best model by AIC in 200 randomly selected microbial SADs**

| Model | Sampling error structure | Best model | Statistically significant * |
|---|---|---|---|
| Lognormal | Poisson | 12.0% | 4.0% |
| | Negative Binomial | 0.5% | 0.0% |
| Logseries | None | 0.5% | 0.0% |
| | Poisson | 0.0% | 0.0% |
| | Negative Binomial | 0.0% | 0.0% |
| Power law | None | 2.5% | 0.0% |
| | Poisson | 13.5% | 0.5% |
| | Negative Binomial | 2.0% | 1.0% |
| Powerbend | None | 5.0% | 2.5% |
| | Poisson | 52.5% | 39.5% |
| | Negative Binomial | 11.5% | 9.5% |

A best model is considered statistically significant when its AIC difference to the second-best model is greater than 2.

**References**


1. Pueyo, S. Diversity: between neutrality and structure. *Oikos* **112**, 392–405 (2006).

2. Fisher, R. A., Corbet, A. S. & Williams, C. B. The relation between the number of species and the number of individuals in a random sample of an animal population. *J Animal Ecol* **12**, 42 (1943).

3. MacArthur, R. H. On the relative abundance of bird species. *Proc. Natl Acad. Sci.* **43**, 293–295 (1957).

4. May, R. M. Patterns of Species Abundance and Diversity. in *Ecology and Evolution of Communities* (eds. Cody, M. L. & Diamond, J. M.) 81–120 (Harvard University Press, 1975).

5. Motomura, I. A statistical treatment of ecological communities. *Zoological Magazine* **44**, 379–383 (1932).
